# Cancer cells survival is dependent on the lincRNA JUNI

**DOI:** 10.1101/2022.06.21.496924

**Authors:** Vikash Kumar, Xavier Sabaté-Cadenas, Isha Soni, Esther Stern, Vias Carine, Doron Ginsberg, Martin Dodel, Faraz K. Mardakheh, Alena Shkumatava, Eitan Shaulian

**Affiliations:** Dep of Biochemistry and Molecular Biology, Institute for Medical Research Israel-Canada, Faculty of Medicine, Hebrew University of Jerusalem, Israel; Institut Curie, PSL Research University, CNRS UMR3215, INSERM U934, Paris 75005, France; The Mina and Everard Goodman, Faculty of Life Science, Bar-Ilan University, Ramat Gan, Israel; Centre for Cancer Cell and Molecular Biology, Barts Cancer Institute, Queen Mary University of London, John Vane Science Centre, Charterhouse Square, London EC1M 6BQ, UK; Gene Therapy Institute, Hadassah Hebrew University Medical Center and Faculty of Medicine, Hebrew University of Jerusalem, Israel

## Abstract

Identification of key factors for cellular survival is a basis for therapy. We identified *JUNI* (linc01135) as a stress-regulated lncRNA, essential for cell survival and implicated in cancer. Besides regulating c-Jun expression we demonstrate c-Jun-independent, robust requirement for cell survival. Analysis of median survival of cancer patients suffering from various types of cancer reveals correlations of *JUNI* expression levels with alterations in patients’ survival. *JUNI’s* antagonistic interaction with DUSP14, a negative regulator of the JNK pathway, underlies the regulation of c-Jun and partial effects on cellular survival. Consistently, DUSP14 expression is coherently inversely-correlated with the survival of patients suffering from the same types of cancer. Our data suggests that *JUNI* is a novel master regulator of cell fate.

**Summary:** JUNI is a novel regulator of cell survival, JNK activation and c-Jun expression, implicated in survival of cancer patients

---

The transcription factor c-Jun is a major integrator of various extracellular or stress signaling which controls cell fate by mediating a wide range of transcriptional responses governing processes such as proliferation, cell death, migration, drug resistance, wound healing, epithelial to mesenchymal transition (EMT) and more(*2-13*). To identify genes regulated by c-Jun, we analyzed ChIP-seq data from ENCODE for genes whose transcription start sites (TSS) are located in the vicinity of c-Jun binding sites. One of the genes whose TSS is located only 1100 bp away from *JUN* TSS is the long intergenic noncoding RNA, linc01135, referred to hereafter as *JUNI* (for *JUN* inducer). According to ENCODE data, *JUNI* contains five main exons and has multiple isoforms. 28 different transcript isoforms were described according to the LNCipedia (*14*). Importantly, the ENCODE predicts that the first exon is shared by all. *JUNI* is evolutionary conserved within primates (Fig S1). The fact that c-Jun is known to bind to its own promoter and to autoregulate its expression (*15*) supported the ChIP-seq results, but did not confirm any functionality for *JUNI*. Therefore, we analyzed its expression to exclude promoter leakiness. UV-driven activation of the c-Jun N-terminal kinase (JNK) results in JNK-dependent c-Jun phosphorylation, transcript elevation and protein stabilization (*16-18*). Hypothesizing that expression of *JUNI* is regulated by c-Jun, and given that UV radiation is a major inducer of c-Jun expression, we irradiated four different cell types with UV radiation and examined *JUNI* expression. In this study we used a set of cell lines including two melanoma cell lines, HMCB and CHL1, HeLa (cervical carcinoma cell line in which *JUN* expression was intensively studied) and MDA-MB-231 (breast cancer-TNBC; a cancer type in which *JUN* plays a significant role) (*19*). Irradiation of these cell lines with 20-30 J/m^2^ UVC resulted in three to five-fold induction of *JUNI* expression (Fig 1 A). Similar to *JUN*, the induction was dose dependent (Fig 1B) and the rapid response to stress (Fig 1C) as well as to serum stimulation of starved cells (*20*), qualifies it as an “immediate early” lncRNA. To determine how common is the correlation between *JUNI* and *JUN* induction following exposure to drug-induced stress, HeLa cells were exposed to chemotherapeutic drugs that induce *JUN* expression and *JUNI* levels were monitored. Every DNA damaging treatment that induced *JUN* also induced *JUNI*, although to a lower extent (Fig 1 D). As *JUN* promoter is the major regulatory element proximate to the first exon of *JUNI* we tested if it co-regulates *JUNI*’s expression. We transfected HeLa and MDA-MB-231 cells with a genomic element that contains the promoter of *JUN* flanked by 153 bases of the first exon of *JUNI* on one side and 750 bp of the 5’ UTR of *JUN* on the other side. Examination of RNA extracted after DNase treatment demonstrated eight and 21-fold higher expression of *JUNI* relatively to cells transfected with an empty vector, in the different cell lines (Fig 1E). Thus, suggesting that *JUN*’s promoter is bidirectional, driving expression of *JUN* on one side and *JUNI* on the other, as previously described for other lncRNAs (*21*). Fractionation experiments demonstrated that *JUNI* resides mainly in the nucleus (Fig. 1F). To define the pathways required for *JUNI* upregulation after stress exposure we inhibited two UV-activated kinases that are known to phosphorylate/induce c-Jun, JNK and p38 using specific inhibitors SP600125 (JNK) or SB203580 (p38). Both inhibitors abolished *JUNI*’s induction by UV similarly to c-Jun expression (Fig S2), suggesting that expression of *JUNI* may be c-Jun-dependent. However, siRNA-directed depletion or overexpression of *JUN* had no significant consequences on *JUNI* expression, suggesting that *JUNI* is not regulated by c-Jun (Fig 1G, H and Fig 4 A, C, E).

**Figure 1.**
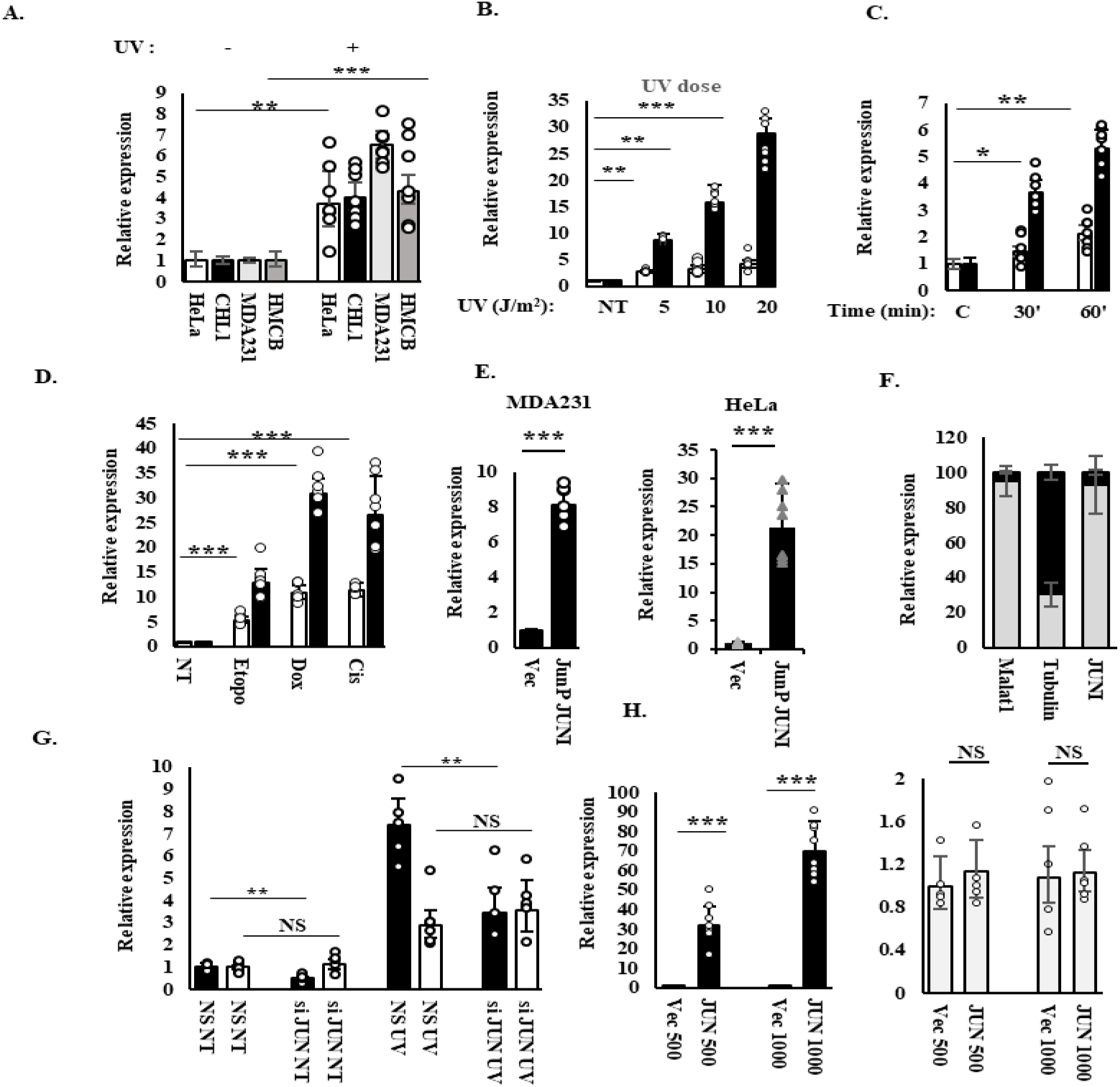
The regulation of *JUNI* A. Expression of *JUNI* in the indicated cell lines before and 4 h after exposure to 20 (HeLa) or 30 J/m^2^ UV (Other indicated cell lines). GAPDH was used for normalization in this experiment and in all the following qPCR experiments. B. HeLa cells were irradiated with the indicated doses of UV and harvested 4 h after irradiation. *JUNI* and *JUN* RNA levels (white and black bars respectively) were determined with RTqPCR. C. HeLa cells were irradiated with 20 J/m^2^ UV and harvested 30 or 60 min after irradiation. *JUNI* and *JUN* RNA levels (white and black bars respectively) were determined with RTqPCR D. HeLa cells were untreated (NT) or exposed to 5μM of Etoposide, doxorubicin or cisplatin. *JUN* and *JUNI* RNA levels (black and white bars respectively) were determined 4h after treatment using RTqPCR. E. MDA-MB-231 and HeLa cells were transfected with a vector containing the genomic sequences of *JUN* promoter and the adjacent first exon of *JUNI* or an empty vector. 48h later *JUNI* expression was monitored using qPCR after DNase treatment. F. HeLa cells were fractionated to nuclear and cytoplasmic fractions and the abundance of *JUNI* was examined in each by RTqPCR. Gray portion of the bar represents the nuclear fraction and black represents the cytoplasmic one. RNAs of Tubulin and MALAT1 were used as markers for cytoplasmic and nuclear fractions respectively G. HeLa cells were transfected with nonspecific (NS) or *c-Jun* siRNAs, irradiated (UV) or not (NT) with 20 J/m^2^ UV and harvested 4 h after irradiation. The levels of JUN RNA (Black bars) or *JUNI* (White bars) were measured by RTqPCR. (upper NS = Non-significant). H. HeLa cells were transfected with the indicated concentrations of *JUN* expression or empty vector (ng), harvested 48h later and levels of *JUN* RNA (Black bars) or *JUNI* (White bars) were measured by RTqPCR. Keys for significance marks; * = p<0.05, ** = p<0.001, *** =p< 1×10^−5^

Because many lncRNA exert their biological effects by modulating the expression of their neighbor genes in *cis* (*22*), we examined the effects of *JUNI* on *JUN* expression. To this end we attempted to knockout the first exon of *JUNI* using CRISPR gene editing targeting *JUNI* with two sgRNAs flanking its first exon. However, exon 1-deleted clones could not be generated, suggesting an essential role for this gene (Fig 3). As the first exon is common to all isoforms we designed two siRNAs targeting it, and used them to transiently silence *JUNI* expression. Both siRNAs silenced *JUNI* and consequently *JUN* mRNA expression, to a lesser degree (Fig. 2A). It is worth mentioning that cell line specific, consistent variations in the silencing capacities of the two siRNAs were observed. Importantly, all the phenotypic effects described in this study corresponded to the silencing efficiencies of each siRNA. HMCB and MDA-MB-231 cells express high levels of c-Jun protein, that can be detected even without stress exposure. Reduction in c-Jun protein levels after *JUNI* silencing was detected in these cells (Fig 2B). In contrast, a clear effect of silencing can be observed in all cell lines after exposure to UV (Fig 2C), suggesting that *JUNI* has special importance for c-Jun expression after exposure of cells to stress. Indeed, exogenous expression of the first exon of *JUNI* either transiently (Fig 2 D) or stably (Fig 2E) was sufficient to elevate c-Jun levels in cells exposed to UV radiation. Combined, our data suggests that *JUNI* regulates *JUN* expression in *trans*, and is particularly important after stress exposure. Significant correlations between *JUNI* and *JUN* expression also occur *in-vivo*. Examination of data from 32 types of tumors in the Pan-Cancer Co-Expression Analysis for the RNA-RNA interactions revealed 94% positive correlation between *JUNI* and *JUN* levels in all cancer types examined. 62% of the positively correlated cases were statistically significant (*23*) (Table S1).

**Figure 2.**
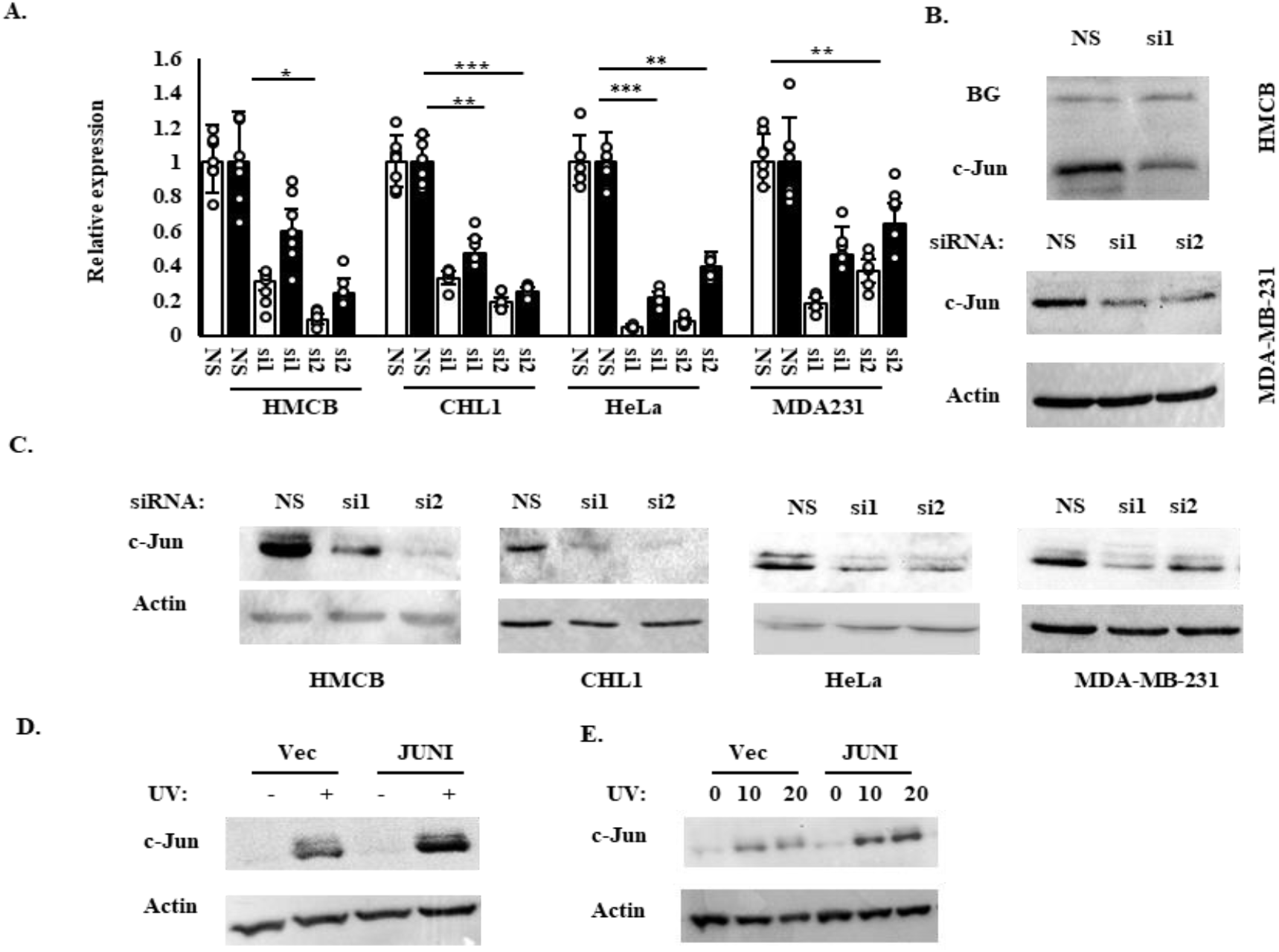
*JUNI* regulates *JUN* expression. A. The indicated cells were transfected with nonspecific siRNA (NS) or two siRNAs against *JUNI* (si1: 20nM and si2: 5 nM). The cells were harvested 24h later and RTqPCR was performed to determine *JUNI* (white bars) and *JUN* (black bars) levels B. HMCB cells (upper part) and MDA-MB-231 cells (lower part) were transfected with NS siRNA or ones against *JUNI* and harvested 48 h later. *c-Jun* protein levels were determined by immunoblotting using specific antibody. Equal background band (BG), or Actin were used as loading control. C. The indicated cells were transfected with NS siRNA or siRNAs against *JUNI* and exposed to UV radiation. *c-Jun* protein levels were examined by immunoblotting 4-6 h after exposure using specific antibody. Actin was used as a loading control. HeLa cells were transiently (D) or stably transfected (E) with the first exon of *JUNI* or with an empty vector. Cells were irradiated 36 h post transfection with 20 J/m^2^ (D) or with the indicated UV doses and the levels of *c-Jun* protein were measured 4 h later as above.

**Figure 3.**
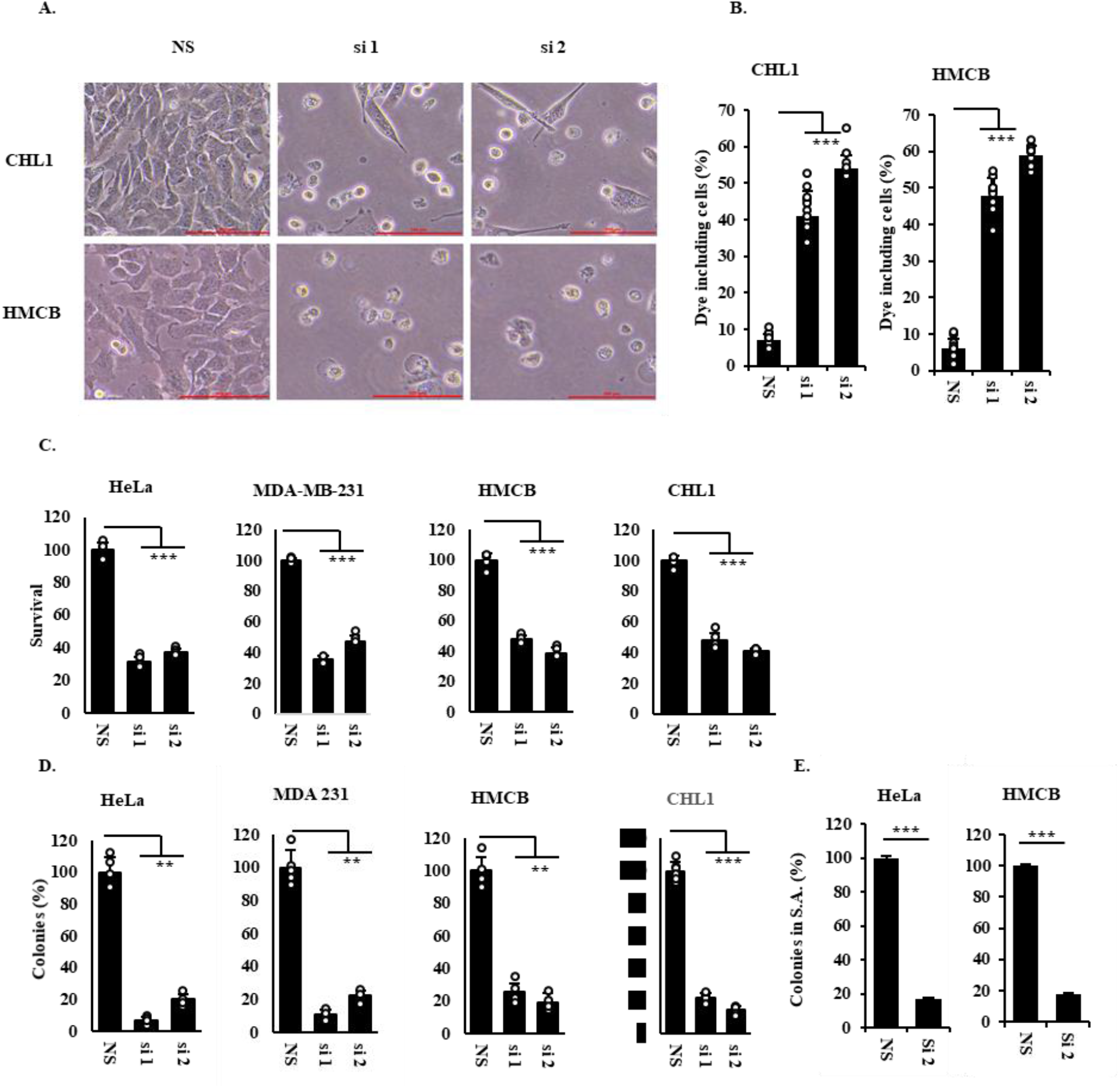
Prolonged silencing of *JUNI* results in cells death. A. The indicated cells were transfected with the indicated siRNAs and photographed 120 h later. B. Trypan blue absorbing cells 96 h post transfection with the indicated siRNA were counted and presented as percent of the total population C. Graphic presentation of the percentage of cellular survival of the indicated cells measured using XTT 120 h post transfections with different siRNAs. Survival of cells transfected with NS siRNA was considered as 100%. D. Clonogenic assay of the indicated cells 14 days after transient transfection with NS, si1 or si2. No selection was used. The number of colonies formed by NS transfected cells was considered as 100%. E. The indicated cell lines were transfected with NS siRNA or *JUNI* siRNA and 24h later cells were plated into soft agar. Clones in 10 fields were counted 2 weeks later. The number of colonies developed in NS transfected cells was considered as 100%.

Next, we examined if *JUNI* has a regulatory effect on targets downstream of c-Jun. As c-Jun can modulate melanoma cells plasticity (*24*) we examined the expression of a set of known c-Jun regulated genes in CHL1 cells 24h after silencing of *JUNI*. Interestingly, c-Jun target genes known to be involved in EMT such as ZEB2 (*25, 26*) and SNAI1 (*27*) were down-regulated by *JUNI* silencing whereas CDH2, which is not a known target, was not (Fig S3 A). The effect on these EMT-relevant, *JUN* targets, was also associated with reduced motility, as reflected by the capacity of cells to migrate in transwell migration assays (Fig. S3 B).

The induction of *JUNI* by DNA damaging agents (Fig 1 A-D) suggested that it may play a role in the DNA damage response. To examine this possibility, *JUNI* was silenced in HMCB, MDA-MB-231 and CHL1 cells which 36-48h later were either UV irradiated or treated with the chemotherapeutic drugs Doxorubicin or Etoposide. Cell survival was determined microscopically and using XTT assays. Silencing of *JUNI* in all cell lines sensitized them to UV-induced-stress and to chemotherapeutic drugs. 30 to 65% reduction in cell survival relatively to cells transfected with control siRNA was observed under the assay condition (Fig. S4). Moreover, we observed apoptotic cells exhibiting blebbing membranes (Fig S4 A).

More striking was the fact that *JUNI* silencing for a prolonged period of 96-120 h in all four examined cell lines resulted in elevated cell death even without exposure to additional stress (Figure 3). Microscopically, the cell death resembled necrotic death with cytoplasmic explosion and free dye entry to the cells (Fig 3A-B), however, some apoptosis and DNA loss was observed as well (Data not shown). Quantification of cell survival using XTT, 120h post *JUNI* silencing revealed 50-70% cell death in the different cell lines (Fig 3C). The survival rate was further reduced to 25-10% two weeks after silencing as indicated by the number of colonies that developed in *JUNI* silenced cells (Fig 3 D). Interestingly, perfect correlation between the capacity of each siRNA to suppress *JUNI* and cell death magnitude, was observed (Fig 2 and Fig 3C-D). Consistently with these results, the silencing of *JUNI* resulted in reduced formation of HeLa and MDA-MB-231 colonies in the soft agar assay (Figure 3E).

As c-Jun is a major cellular signaling molecule involved in many aspects of cellular wellbeing the reduction in cellular survival can potentially be caused by its repression after *JUNI* silencing. To address this possibility, we compared the effects inflicted on survival after *JUNI* silencing to those observed after *JUN* silencing. We used specific siRNAs to silence *JUNI* or *JUN* in 3 different cell lines MDA-MB-231, HeLa and HMCB (Figure 4 A-F). Interestingly, despite of the fact that *JUN* mRNA levels in cells transfected with specific *JUN* siRNA were at least 20% lower than these in *JUNI* silenced cells (Fig 4 A,C,E left panels), their survival 96h after *JUN* silencing was 15-30% higher (Fig 4 A,C,E right panels). This difference extenuated two weeks later. A reduction of 45-60% in colonies formation in *JUNI* silenced relative to *JUN* silenced cells was observed (Fig 4 B, D, F). These results demonstrate that in the examined cell lines *JUNI* is significantly more important for cellular survival than *JUN*. As we demonstrated (Fig. 1 G, H), *JUN* silencing had no significant effect on *JUNI* expression (Fig 4 A,C,E left panels).

**Figure 4.**
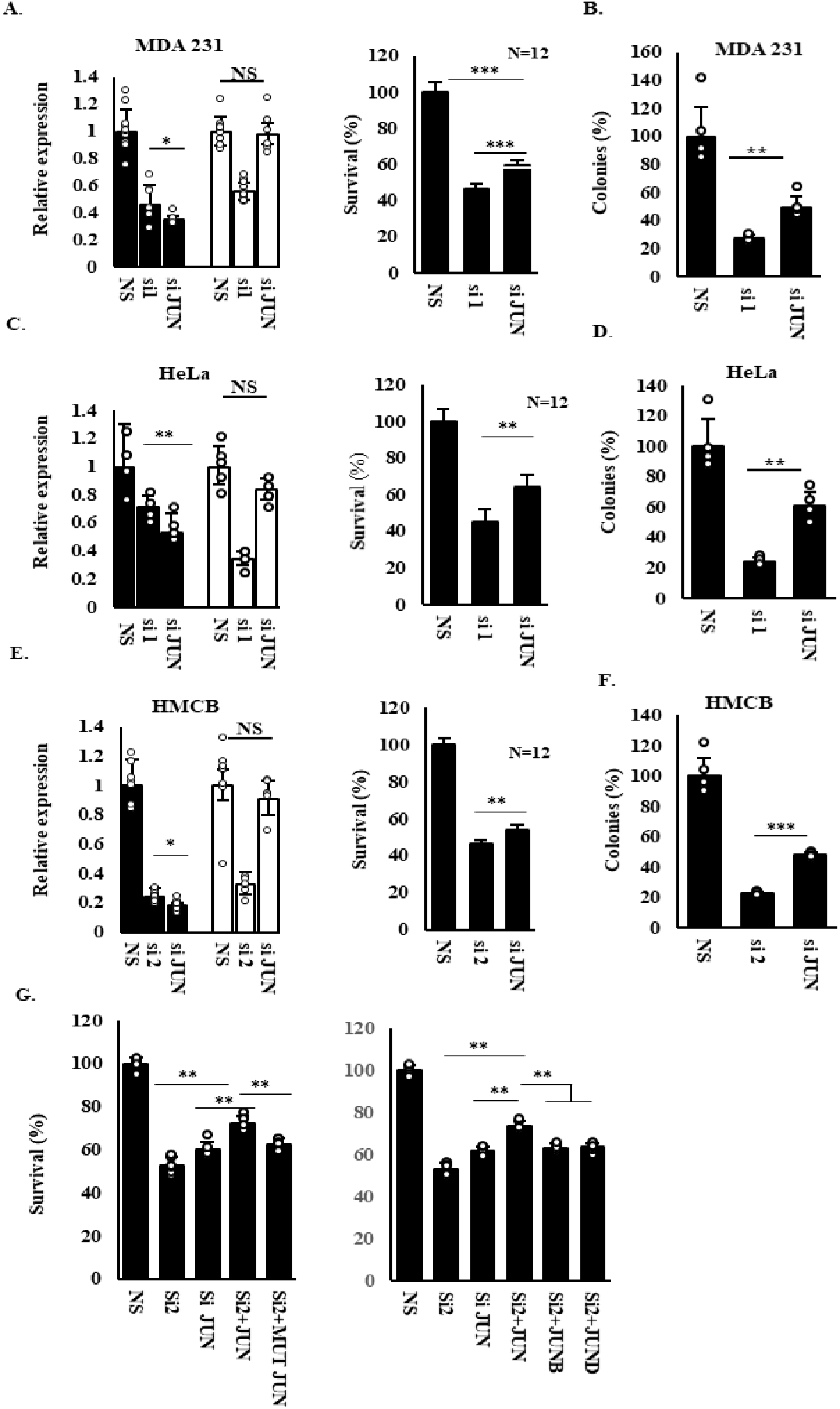
c-Jun mediates only part of *JUNI’*s effects on survival. MDA-MB-231 (A-B) HeLa (C-D) and HMCB cells (E-F) were transfected with NS, siRNA against *JUNI* or siRNA against *JUN*. The levels of *JUN* (black bars) and *JUNI* (white bars) were determined 30h after transfection using RTqPCR (A,C,E left panel). Viability of the transfected cells was determined using XTT 96 h or 120 h later in HeLa or MDA-MB-231 and HMCB respectively (A,C,E right panel), or by counting colonies formed two weeks post transfection (B,D,E). G. HMCB cells were silenced for *JUNI* and co-transfected with the indicated expression vectors. Cell survival was examined using XTT assay 96h post transfection

To further test the effect of c-Jun on *JUNI*’s survival activity, we performed rescue experiments in which *JUNI* was silenced in HMCB cells and exogenous c-Jun was expressed to complement for the repressed endogenous gene. *JUNI* silencing reduced cells survival by 50% whereas *JUN* silencing reduced it by 40% (Fig. 4G). Restoration of *JUN* expression enhanced survival of *JUNI* silenced cells. However, the enhancement was partial (Fig. 4G). Mutant c-Jun that cannot bind DNA (272/273E) (*28*) or other *JUN* family members, *JUNB* or *JUND*, could not rescue from *JUNI*-silencing-dependent cell death, thus, suggesting specificity of the c-Jun rescue. These results indicate that *JUNI*s effects on cellular survival are only partially dependent on *JUN* expression and predicts that it may regulate the activity of other factors essential for cellular survival.

To explore the identity of protein interactors that may affect the ability of *JUNI* to regulate c-Jun expression and affect cell death we identified protein interaction partners of *JUNI* by applying incPRINT screen (*26*). Using the shortest isoform of *JUNI* that contained the *JUN* affecting sequences as RNA bait, we identified 57 *JUNI* interacting proteins (Fig. S5). Analyses of their cellular functions revealed enrichment of proteins involved in various fundamental, basic aspects of cellular wellbeing. Dual specificity protein phosphatase 14 (DUSP14; also known as MKP6) was one of the highly rated interacting proteins with *JUNI* (Fig S5 and Fig 5A). DUSP14 can directly dephosphorylate or indirectly limit phosphorylation of MAPKs essential for c-Jun expression JNK, p38 and ERK (*1*). We validated the interaction between DUSP14 and *JUNI* interaction in HeLa cells by Crosslinking and Immunoprecipitation (CLIP). We examined the enrichment of endogenous *JUNI* following immunoprecipitation with transfected DUSP14 or GFP. Specificity was determined by comparison of *JUNI* enrichment to other unrelated nuclear lncRNAs *MALAT1* and *PVT1*. These experiments demonstrated specific association of *JUNI* with DUSP14 (Fig. 5B). Furthermore, JUNI’s interactions with DUSP14 were significantly stronger after UV exposure of HMCB cells (Data not shown). As *JUNI* is essential for efficient c-Jun upregulation after UV exposure and DUSP14’s inhibits JNK which upregulates *JUN* transcription and protein expression (*16, 17*) we sought to test if *JUNI* antagonizes DUSP14 activity to enable full scale c-Jun induction post stress. Hence, DUSP14 co-silencing with *JUNI* will restore JNK activity, c-Jun phosphorylation and expression in UV irradiated cells. To examine this hypothesis, we silenced DUSP14, *JUNI* or both in HMCB cells that were later exposed to UV radiation, and JNK activation, c-Jun phosphorylation and level were determined. DUSP14 silencing elevated JNK phosphorylation (pThr183/pTyr185) in UV treated cells and consequently Ser 63 phosphorylation of c-Jun and its total level (Fig. 5C). In addition, silencing of *JUNI* inhibited JNK phosphorylation and the consequent c-Jun phosphorylation and induction. JNK phosphorylation was reduced and hardly detected even after exposure to UV irradiation, which is the strongest JNK activator. Co-silencing of DUSP14 together with *JUNI* restored JNK and c-Jun phosphorylation to the levels comparable to non-silenced cells. Out data suggest that *JUNI* regulates JNK phosphorylation and c-Jun’s levels by antagonizing DUSP14. We next examined if antagonism between *JUNI* and DUSP14 also affects biological properties. To this end *JUNI*, DUSP14, or both simultaneously were silenced in HMCB and HeLa that were later treated with UV. In this experiment, cell survival was measured 10h post irradiation. Consistent with our other experiments (Fig. S4), *JUNI* silenced cells are 40% more sensitive to radiation than non-silenced cells (Fig. 5D). DUSP14 silencing did not significantly increase survival of irradiated cells in which *JUNI* expression was normal. Remarkably, it did increase the survival of *JUNI* silenced cells by 29 and 33% in HMCB and HeLa cells respectively (Fig. 5D). However, the negative effect of *JUNI* silencing on survival was not fully rescued. This result suggests *JUNI*-DUSP14 antagonism in regulation of survival of UV exposed cells. Next, we examined whether *JUNI* has any relevance in human cancer and whether the antagonism between *JUNI* and DUSP14 occurs also *in vivo*. We examined the correlation between expression of *JUNI* and survival of patients diagnosed with one of the 21 types of cancer for which RNA seq data is available in the Pan-cancer Project (*29*). In 10 out of the 21 types of cancer examined, we found statistically significant correlations between *JUNI* expression and either decreased or improved patients’ survival, dependent on the cancer type (Table S2). Moreover, the effects of DUSP14 on the median survival of patients in eight out of these 10 cancers was strictly inverse (Fig. 5E). These results suggest that *JUNI* has context-dependent roles in human cancer and that, similar to cell lines, it antagonizes DUSP14 also, in cancer. Notably, unlike DUSP14, expression of *JUN* itself does not negatively or positively correlate with *JUNI* in the prognosis of cancer patients (Data not shown).

**Figure 5.**
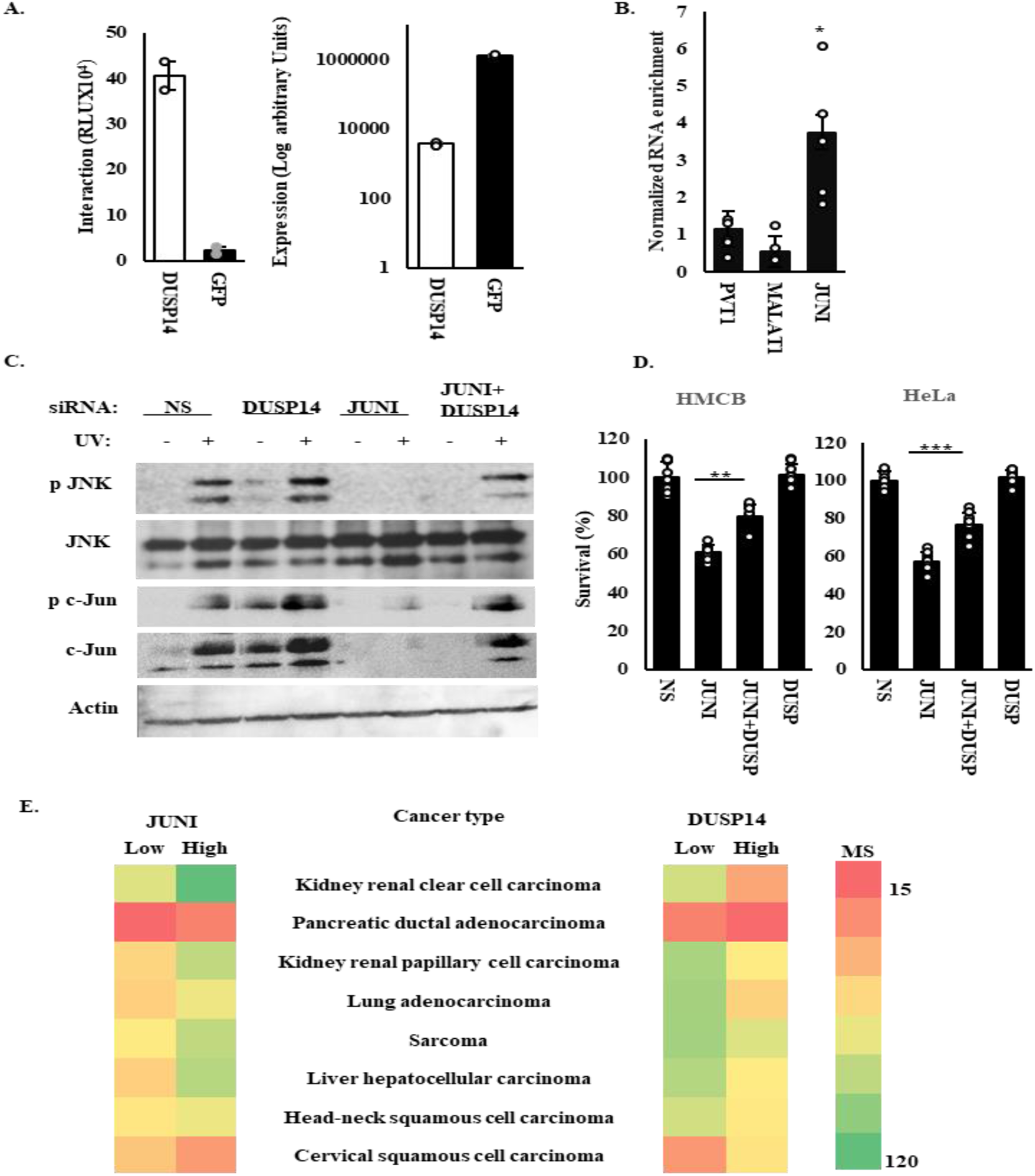
*JUNI* interacts and antagonizes DUSP14 in cells and in cancer A. Interaction between *JUNI* and DUSP14 measured by incPRINT. Left panel, interaction intensities between *JUNI* and the indicated proteins measured by luminesces emissions. Right panel, expression levels of each protein measured by ELISA. GFP was used as non-binder control. Data from two biological replicates are presented as mean ± s.d. RLU are relative light units. B. HeLa cells were transfected with DUSP14 or GFP. UV-crosslinking-immunoprecipitation experiments were performed. The ratio between RNAs co-precipitated with DUSP14 and RNAs in the whole cell extract were further normalized to the ratio obtained from GFP precipitation. Enrichment of the indicated lncRNAs is depicted. C. HMCB cells were transfected with the indicated siRNAs, irradiated with 30 J/m^2^ 48 h later and harvested 5 h post irradiation. Levels of the indicated proteins were measured by immunoblotting using specific antibodies as described before. D. The indicated cells were transfected with the indicated siRNAs, UV irradiated 40 h later and harvested 12 h after irradiation to determine viability using XTT. Survival of NS transfected cells was considered 100%. E. Correlation of the median survival (months) of patients suffering from the indicated types of cancer and expression levels of the indicated genes. In the right, color scale for the time representation in 15 months intervals. Results in the last two types of cancer are specific for a white population.

Taken together, we discovered a novel lncRNA which plays fundamental roles in cellular survival. Aside from regulating the major JNK-JUN cellular pathway, *JUNI* elicits c-Jun-independent, significant effects on cellular survival. Sensitization of the cell lines examined in this study to chemotherapeutic drugs occurs prior to the spontaneous cell death observed at least 48 h later. Given the different kinetics and morphology of cell death, we cannot exclude different cell death mechanisms. The plethora of functions of the putatively interacting proteins may suggest that prolonged silencing leads to a synthetic cell death. *JUNI*-interacting DUSP14 inhibits stress - induced JNK activity and c-Jun induction. As JNK activity induces cell death post UV exposure (*30*) the ability of DUSP14 silencing to rescue from it is surprising. Two potential mechanisms may account for the rescue. First, DUSP14 silencing increase the activity of protective proteins which are negatively regulated by it. For example, NFкB is downstream to the DUSP14 negatively regulated TAK1-TAB1 proteins (*31*). Second, low constitutive JNK activation may induce autophagy and exerts a protective activity (*32, 33*). Sporadic references to *JUNI* as a cancer-relevant, poorly characterized lncRNA have been recently reported (*34, 35*). How *JUNI* impacts tumorigenesis and consequently survival of cancer patients is a ground opened for research.

## Supporting information

Kumar et al Bi...v 21.6.22.pdf

## Acknowledgments

We thank Dr. M. Saleh for the His-DUSP14 plasmid. We thank Ehud Razin’s laboratory for assistance with various reagents.

## Author contributions

Conceptualization: ES, VK

Intellectual input: ES, AS, DG

incPRINT establishment and analysis: AS, XSC, VC

CLIP analysis: FKM, MD

Investigation: VK, XSC, IS, ES, VC, MD,

Supervision: ES, AS FKM

Writing – original draft: ES

Writing – review & editing: ES, AS DG

