## Supplementary material for "Cancer cells survival is dependent on the lincRNA JUNI": Kumar et al Bi...v 21.6.22.pdf

### Supplementary Materials

#### Materials and Methods

##### Cell culture

CHL-1, Hela, MDA-MB 231 and HMCB cells were procured from ATCC. The first 3 were cultured in DMEM with 10% FBS whereas HMCB was cultured in EMEM. All cells were grown with 10% FBS and 1% penicillin-streptomycin cultured at 37°C with 5% CO2. Cells were routinely monitored for mycoplasma contamination.

##### Transfection, treatments, and Inhibitors

Plasmid transfections were carried out with PloyJet reagent (SignaGen Lab MD, USA) according to the manufacturer instructions using 40-50% confluent cells. siRNA transfections were carried out with siRNA transfection reagent: INTERFERin® (Polyplus, Illkirch – France). siRNAs concentrations used in this study ranged between 5 to 20nm. 10µM of SP600125, or SB203580 JNK and p38 inhibitors, were used to inhibit the kinases. Inhibitors were added one hour prior to UV irradiation. Doxorubicin, etoposide and cisplatin were used at .5 to 5 µM dose. XTT reagent was purchased from Abcam (Abcam, Cambridge UK) and used according to the manufacturer instructions.

##### RNA Extraction and Real-Time PCR

RNA was isolated manually using TRI Reagent® (MRC, OH, USA). cDNA was prepared from total RNA (1µg), using Quantabio, qScript™ cDNA synthesis kit and PerfeCTa SYBR Green SuperMix was used for qPCR according to manufacturer's instruction (Quantabio, MA, USA). To prepare DNA-free RNA samples, RNA was treated with PerfeCTa® DNase I according to the manufacturer instructions (Quantabio, MA, USA). StepOnePlus Real-Time PCR apparatus with StepOne Software v2.3 was used for analysis (Applied Biosystems, MA USA). Each experiment was performed in multiple experimental duplications n>4 and least 3 biological repeats. Representative results are depicted

##### Western Blot and Antibodies

Primary antibodies used were c-Jun (CST #60A8 1:1000), Ser-63-p-JUN (CST #9261 1:500), Actin (CST #3700S 1:5000), GAPDH (CST #97166 1:5000), P-JNK (CST #9251 1:500), total JNK total (#9252, #9258 1:1000), anti-FLAG M2 antibody (Sigma, #F1804). HRP-conjugated secondary antibody was used at a dilution of 1:5000 (Jackson Laboratories, USA).

##### Colony formation and soft agar assays

For colony formation assay 1000-2000 cells were seeded in 3.5 cm dishes and cultured for 12-14 days. The cells were fixed using Methanol: Acetic acid (3:1) ratio. Colonies were stained with Methylene Blue using 1% methylene blue in 0.1M borate buffer, pH 8.5. For soft agar assays 10k-15k cells were seeded on 1%

agar/complete media mixture coated 3.5cm plates, covered with 0.3% agar/complete media mixture and cells were incubated for 15-20 days.

**Microscopy**

Images were captured at 20X using Inverted NIKON ECLIPSE T2S microscope. 50 μm Scale bar are presented.

**Plasmids and cloning**

Plasmids expressing *JUNB*, *JUND* and Mutant *JUN* that cannot bind DNA (272/273E) were previously described (*28*). DUSP14-HIS was a kind gift of M. Saleh (*36*). To generate *JUNI*-MS2X10 construct used in the incPRINT, 938 bp fragment of the *JUNI* mature RNA was cloned into 10XMS2 vector (*26*) using a gBlocks Gene Fragment containing Cla1 and Hind3 restriction sites (IDT, NJ, USA). Exon1 of *JUNI* was cloned into pEFA1 CMV puro/GFP vector using a gBlocks gene fragment containing EcoR1 and BAMH1 restriction enzymes.

**Crosslinking and Immunoprecipitation (CLIP):**

10 million HeLa cells were transiently transfected with mammalian expression vectors expressing His-DUSP14 or GFP as a control, using Lipofectamine 2000 (Thermo Fisher) according to manufacturer's instructions. The next day, cells were washed with PBS and irradiated once on ice with 150 $mJ/cm^2$ UV light (254 nm) in ice-cold PBS, using a Hoefer Scientific UV Crosslinker. Cells were then pelleted and lysed in 1 ml of lysis buffer (50mM Tris-HCl pH 7.4, 100mM NaCl, 1% Igepal CA-630, 0.1% SDS, 0.5% sodium deoxycholate, supplemented with protease inhibitors), sonicated, cleared, and adjusted to a protein concentration of 1mg/ml. RNA was then partially digested with 0.2 U/ml of RNase I for 3 mins, before immunoprecipitation with 10 μL of anti-His-tag antibody (Cell Signaling), pre-conjugated to 50 μL protein G Dynabeads (Thermo Fisher) for 1 hour. The beads were then washed 5 times with lysis buffer, before the bound RNAs were eluted off by Proteinase K (Thermo Fisher) digestion for 1hour at 65°C. The RNA was then extracted using TRIZol (Thermo Fisher), and subjected to qPCR analysis with qPCR primers against *JUNI*, *MALAT1*, and *PVT1*.

**incPRINT**.

384-well plates (Greiner Bio-One, #781074) were coated overnight with the anti-FLAG M2 antibody (Sigma, #F1804; 10 μg/ml in PBS) and blocked for two hours in 1% BSA, 5% sucrose, 0.5% Tween-20 in 1×PBS. Plasmids encoding *JUNI*-10xMS2 and 3×FLAG-tagged test proteins were co-transfected using

Polyethylenimine (PEI) (MW 40,000, Polysciences, #24765) into a HEK 293T stable cell line expressing the NanoLuc luciferase fused to MS2CP (Graindorge et al 2019). The day before transfection, cells were seeded in 96-well plates (30,000 cells per well). Co-transfections were performed using 300 ng of RNA plasmid and 150ng of each protein plasmid per well. Two days after transfection, cells were washed twice with ice-cold 1x PBS and lysed in ice-cold RQ1-HENG buffer (20 mM HEPES-KOH pH 7.9, 150 mM NaCl, 10 mM MgCl2, 1 mM CaCl2, 5% glycerol, 1% Triton X-100), supplemented with protease inhibitors (aprotinin, leupeptin and pepstatin, 1 μg/ml each, PMSF 0.5 mM), 40 U/ml of RNaseOUT Ribonuclease Inhibitor (Invitrogen, #10777-019) and 30 U/ml of RQ1 DNase (Promega, #M6101). After lysis (10 min, 4 °C) and DNase incubation (30 min, 37 °C), the lysates were transferred to 384-well plates previously coated. Following a three-hour incubation at 4 °C, plates were washed seven times with RQ1-HENG buffer. A 1:200 dilution of furimazine substrate (Promega, #N1110) was added to the plates and luminescence in each well was measured with a plate reader (TECAN Spark). Following luminescence measurement, HRP-conjugated anti-FLAG antibody (Abcam, #ab1238, 1:10,000 dilution) in ELISA buffer (1× PBS, 1% goat serum, 1% Tween-20) was added to each well. After 30 min of incubation at room temperature, plates were washed with 1× PBS, 0.05% Tween-20, SuperSignal™ ELISA Pico Chemiluminescent Substrate (Thermo Fisher Scientific, #37069) were added and ELISA signals were detected with the microplate reader. All 96- and 384-well plate washes were performed with a microplate washer (Biotek 405LSUVS). Data was normalized as previously described (*26*).

**Primers and siRNAs**

| PCR primers | | |
|---|---|---|
| **Gene Name** | **Forward** | **Reverse** |
| GAPDH | TCGACAGTCAGCCGCATCTTCTTT | ACCAAATCCGTTGACTCCGACCTT |
| JUN | CCCCAAGATCCTGAAACAGA | CCGTTGCTGGACTGGATTAT |
| JUNI | CGGACACTCGCATAAAGTCA | AGCGCTTCCTAGAGGCTACC |
| Zeb2 | CAAGAGGCGCAAACAAGCC | GGTTGGCAATACCGTCATCC |
| SNAI1 | CTGGGTGCCCTCAAGATGCA | CCGGACATGGCCTTGTAGCA |
| DUSP14 | GAGGCGTACAACTGGGTGAA | CCCCAGTAAGGCATCAGGT |
| CDH2 | TGTCGGTGACAAAGCCCCTG | AGGGCATTGGGATCGTCAGC |
| MALAT1 | GACGGAGGTTGAGATGAAGC | ATTCGGGGCTCTGTAGTCCT |
| PVT1 | TGGAATGTAAGACCCCGACTCT | GATGGCTGTATGTGCCAAGGT |

| siRNAs | |
|---|---|
| JUNI si1 | GACACUCGCAUAAAGUCACGCAGUA |
| JUNI si2 | GGUAGAACCUGGACUUCAAAUCUCU |
| JUN | ACUCAUGCUAACGCAGCAGUU |
| DUSP14 | GGCAUCACCUGCAUUGUUAAUGCTA |

#### Figures

#### Fig. S1.

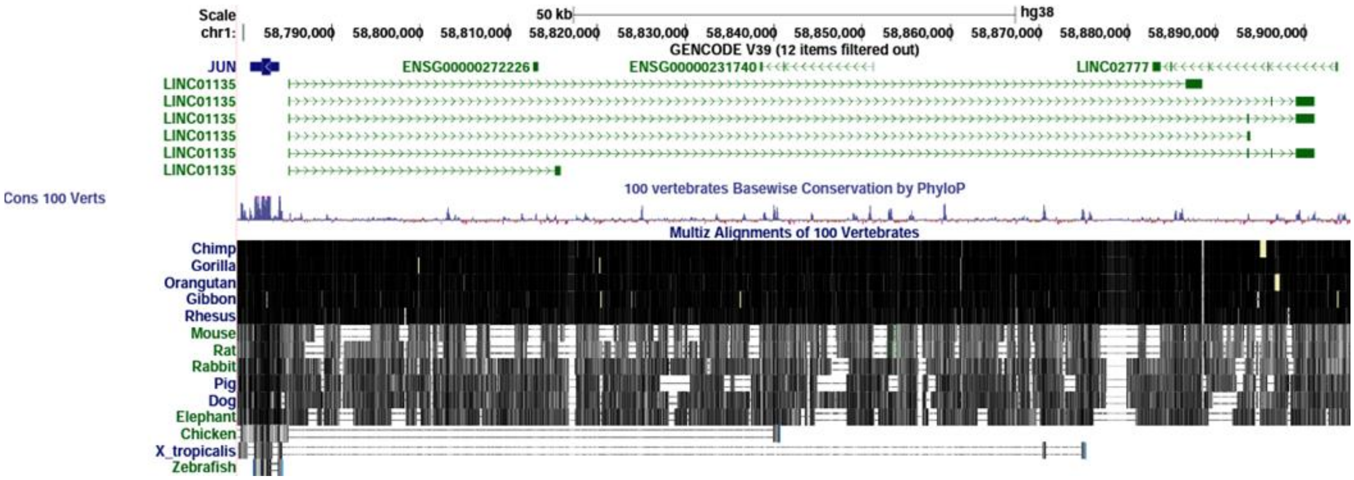


**Evolutionary conservation of *JUNI*.**

Data obtained from UCSC Genome Browser on Human (GRCh38/hg38).

**Fig. S2.**

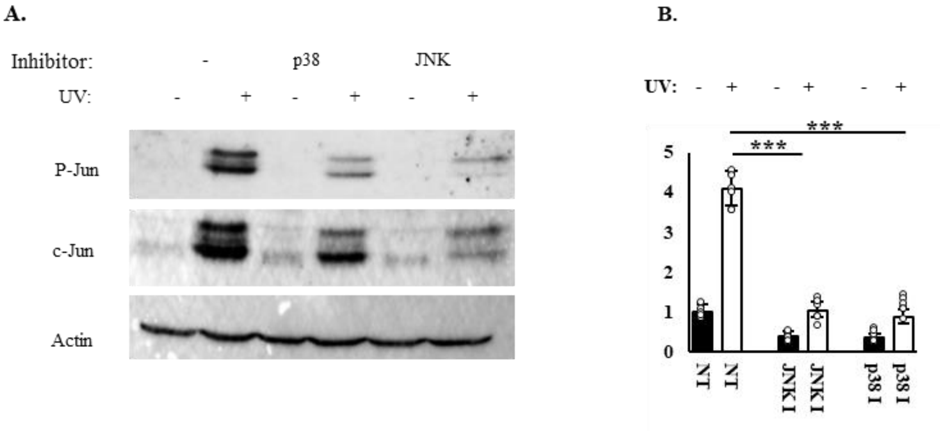


**p38 and JNK regulate *JUNI* expression.**

A. HeLa cells were treated with 10 μM SP600125 (JNK inhibitor) or SB203580 (p38 inhibitor) 1 h prior to exposure to 20 J/m$^2$ UV, and harvested 2 h after exposure. Phosphorylated (Ser 63) and total c-Jun protein levels were determined using specific antibodies. Actin was used as loading control. B. *JUNI* RNA levels were determined with RTqPCR in cells from the same experiment as in A.

**Fig. S3.**

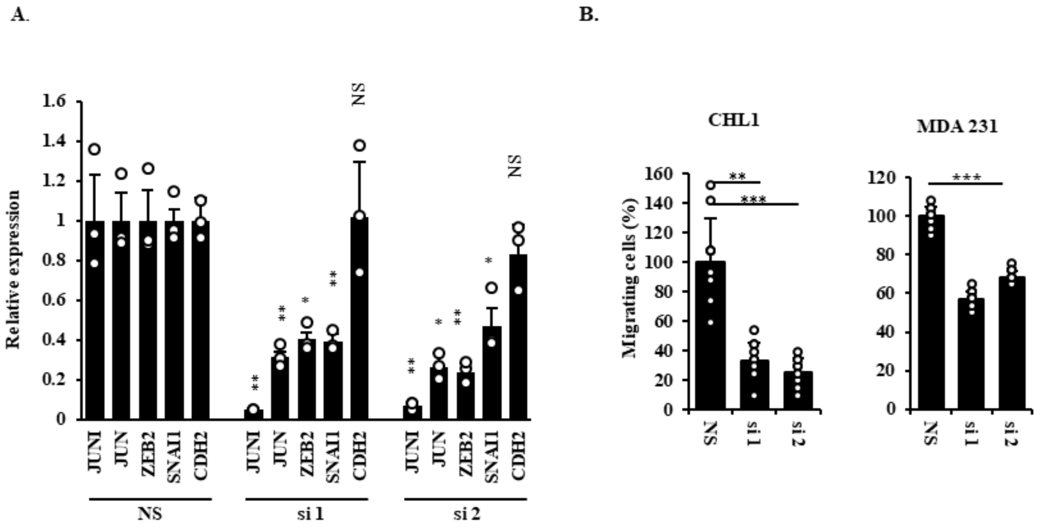


***JUNI* regulates c-Jun targets and cellular motility**

A. CHL1 cells were transfected with the indicated siRNAs. Expression of the indicated genes was determined 24 h later using RTqPCR and specific primers. B. The percentage of siRNA transfected cells that crossed 8-micron millicell hanging inserts membrane within 16 hours post plating. 500000 cells were seeded 24h after transfection with the indicated siRNA. The migration of NS siRNA transfected cells was considered as 100%.

**Fig. S4**

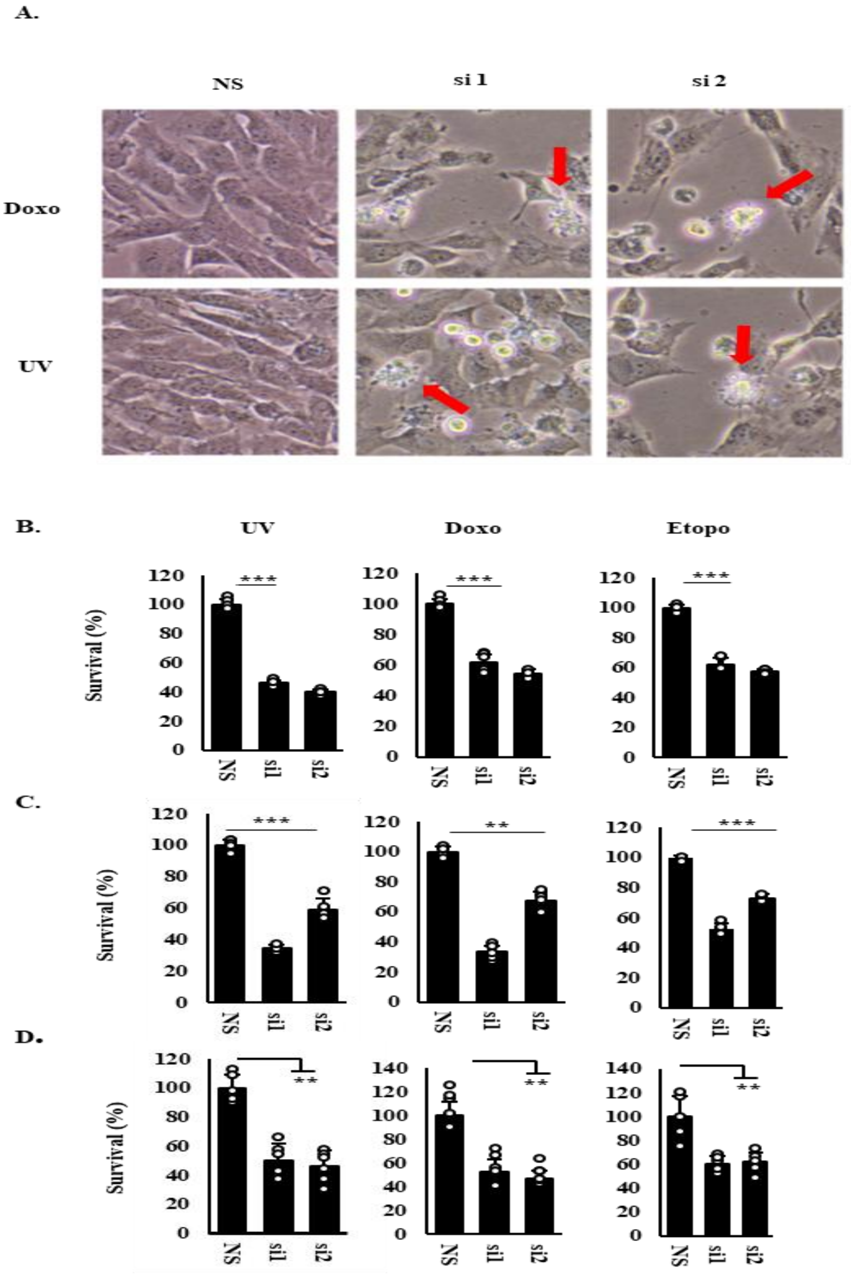


**Silencing of *JUNI* sensitizes cancer cells to chemotherapy-induced cell death.**

A. CHL1 cells were transfected with the indicated siRNAs and 40 h later were irradiated with 30 J/m$^2$ UV or treated with 5 μM doxorubicin and photographed 3 or 5 h later respectively. Characteristic morphologies of apoptotic cells were indicated by red arrows. B. HMCB cells were transfected with the indicated siRNAs and treated 36 h later with 15 J/m$^2$ UV, 1μM doxorubicin or 5μM etoposide. XTT was measured 20 h later. C. MDA-MB-231 cells were transfected with the indicated siRNA and 48 h later were treated with 10 J/m$^2$ UV, 1μM doxorubicin or 5μM etoposide, harvested 12h after UV and 20h after drugs treatment and XTT levels were measured as above. D. CHL1 cells transfected with the indicated siRNA were treated 38 h later with 15 J/m$^2$ UV, 1μM doxorubicin or 5μM etoposide. XTT was measured 20 h later.

**Fig. S5**

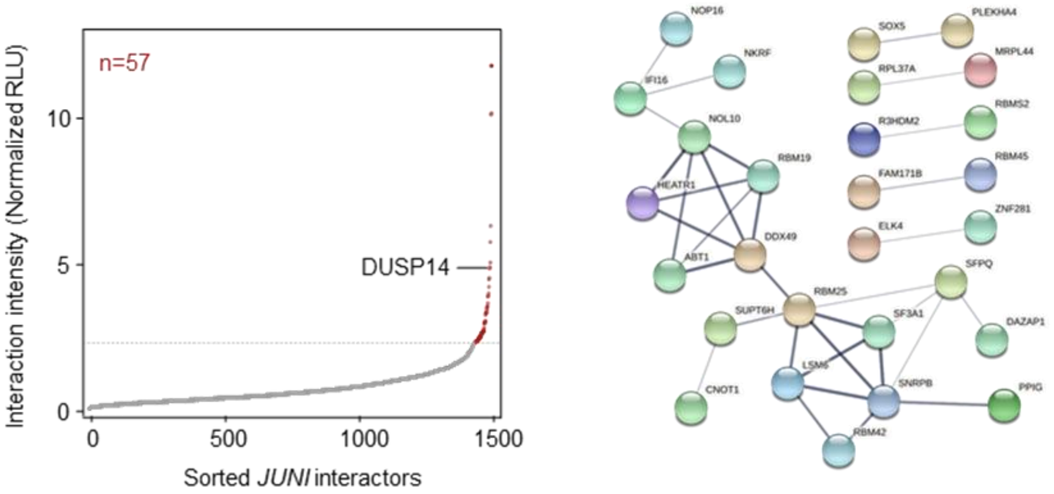


**Comprehensive protein interactome of *JUNI* in cells.**

A. Normalized *JUNI*-protein interaction intensities averaged from two biological replicates, sorted in increasing order. The horizontal dotted line represents interaction intensity cutoff used for the classification of *JUNI* interactors. Red dots are *JUNI*-interacting proteins; gray dots are proteins that do not bind to *JUNI*. See the ‘Methods’ section for data normalization. RLU are relative light units. B. STRING analysis of the 57 *JUNI* interactors. Shown are only connected nodes. Evidence comes from protein-protein interaction sources of experiments, databases, and text-mining. The line thickness indicates the strength of data support: medium confidence (0.4, thin lines), high confidence (0.7, medium lines) or highest confidence (0.9,thick lines). The PPI enrichment p-value <10-5.

**Table S1.**

| **Cancer type** | **Cofficient-R** | **sample size** | **p-value** |
|---|---|---|---|
| Breast Invasive carcinoma | 0.121 | 1104 | **5.20E-05** |
| Cervical Squamous Cell Carcinoma and Endocervical Adenocarcinoma | 0.157 | 306 | **6.02E-03** |
| Adrenocortical carcinoma | 0.187 | 79 | **9.97E-02** |
| Bladder urothelial carcinoma | 0.172 | 411 | **4.69E-04** |
| Cholangiocarcinoma | 0.264 | 36 | **1.20E-01** |
| Colon Adenocarcinoma | 0.033 | 471 | **4.69E-01** |
| Lymphoid Neoplasm Diffuse Large B-cell Lymphoma | 0.415 | 48 | **3.37E-03** |
| Esophageal Carcinoma | 0.277 | 162 | **3.58E-04** |
| Head and Neck Squamous Cell Carcinoma | 0.146 | 502 | **1.02E-03** |
| Kidney Chromophobe | 0.198 | 65 | **1.14E-01** |
| Kidney Renal Clear Cell Carcinoma | 0.094 | 535 | **2.98E-02** |
| Acute Myeloid Leukemia | 0.424 | 151 | **5.94E-08** |
| Brain Lower Grade Glioma | 0.131 | 529 | **2.56E-03** |
| Liver Hepatocellular Carcinoma | 0.237 | 374 | **3.67E-06** |
| Lung Adenocarcinoma | 0.213 | 526 | **7.88E-07** |
| Lung Squamous Cell Carcinoma | 0.162 | 501 | **2.61E-04** |
| Ovarian Serous Cystadenocarcinoma | 0.222 | 379 | **1.30E-05** |
| Pancreatic Adenocarcinoma | 0.239 | 178 | **1.25E-03** |
| Pheochromocytoma and Paraganglioma | 0.338 | 183 | **8.24E-02** |
| Prostate Adenocarcinoma | 0.219 | 499 | **7.71E-01** |
| Rectum Adenocarcinoma | 0.231 | 167 | **2.17E-01** |
| Sarcoma | 0.308 | 263 | **2.87E-05** |
| Skin Cutaneous Melanoma | 0.123 | 471 | **3.02E-02** |
| Stomach Adenocarcinoma | 0.253 | 375 | **1.10E-01** |
| Testicular Germ Cell Tumors | 0.402 | 156 | **2.02E-04** |
| Thyroid Carcinoma | 0.261 | 510 | **1.49E-01** |
| Thymoma | 0.532 | 119 | **1.30E-12** |
| Uterine Corpus Endometrial Carcinoma | 0.136 | 548 | **5.04E-18** |
| Uterine Carcinosarcoma | 0.25 | 56 | **1.23E-01** |
| Uveal Melanoma | 0.285 | 80 | **7.33E-01** |
| Mesothelioma | -0.184 | 86 | **9.07E-02** |

Correlations of *JUNI* and *JUN* co-expression in 32 Cancer types. Data obtained from the Pan-Cancer Co-Expression Analysis for the RNA-RNA interactions (*23*).

**Table S2.**

| **Cancer type** | **Population** | **Survival** | **p-value** | **M.S. low** | **M.S. high** |
|---|---|---|---|---|---|
| Esophageal Squamous Cell Carcinoma | General | Reduced | 0.023 | 48.6 | 21.67 |
| Kidney renal clear cell carcinoma | General | Improved | 5.7 e-05 | 63.73 | 118.47 |
| Kidney renal papillary cell carcinoma | General | Improved | 0.0021 | 43.5 | 77.7 |
| Liver hepatocellular carcinoma | General | Improved | 0.039 | 42.37 | 81.87 |
| Lung adenocarcinoma | General | Improved | 0.015 | 42.17 | 57.5 |
| Lung squamous cell carcinoma | General | Improved | 0.019 | 44.6 | 69.5 |
| Pancreatic ductal adenocarcinoma | General | Improved | 0.0016 | 15.33 | 22.03 |
| Sarcoma | General | Improved | 0.033 | 49.27 | 77.47 |
| Cervical squamous cell carcinoma | Whites | Reduced | 0.05 | 39.53 | 28.7 |
| Head-neck squamous cell carcinoma | Whites | Improved | 0.0489 | 48.63 | 58.27 |

*JUNI*'s expression levels correlates significantly with differences in the survival of cancer patients. Results present analysis of survival of patients expressing high or low levels of *JUNI* using a Kaplan-Meier Plotter (*29*) mediated processing of Pan-cancer RNA-seq data. M.S. = Median Survival in months.
